# Genomes from uncultivated prokaryotes: a comparison of metagenome-assembled and single-amplified genomes

**DOI:** 10.1101/219295

**Authors:** Johannes Alneberg, Christofer M. G. Karlsson, Anna-Maria Divne, Claudia Bergin, Felix Homa, Markus V. Lindh, Luisa W. Hugerth, Thijs JG. Ettema, Stefan Bertilsson, Anders F. Andersson, Jarone Pinhassi

## Abstract

**Background:** Prokaryotes dominate the biosphere and regulate biogeochemical processes essential to all life. Yet, our knowledge about their biology is for the most part limited to the minority that has been successfully cultured. Molecular techniques now allow for obtaining genome sequences of uncultivated prokaryotic taxa, facilitating in-depth analyses that may ultimately improve our understanding of these key organisms.

**Results:** We compared results from two culture-independent strategies for recovering bacterial genomes: single-amplified genomes and metagenome-assembled genomes. Single-amplified genomes were obtained from samples collected at an offshore station in the Baltic Sea Proper and compared to previously obtained metagenome-assembled genomes from a time series at the same station. Among 16 single-amplified genomes analyzed, seven were found to match metagenome-assembled genomes, affiliated with a diverse set of taxa. Notably, genome pairs between the two approaches were nearly identical (>98.7% identity) across overlapping regions (30-80% of each genome). Within matching pairs, the single-amplified genomes were consistently smaller and less complete, whereas the genetic functional profiles were maintained. For the metagenome-assembled genomes, only on average 3.6% of the bases were estimated to be missing from the genomes due to wrongly binned contigs; the metagenome assembly was found to cause incompleteness to a higher degree than the binning procedure.

**Conclusions:** The strong agreement between the single-amplified and metagenome-assembled genomes emphasizes that both methods generate accurate genome information from uncultivated bacteria. Importantly, this implies that the research questions and the available resources are allowed to determine the selection of genomics approach for microbiome studies.

## Background

The genome is a fundamental resource for understanding the physiology, ecology and evolution of an organism. With the availability of high-throughput sequencing technologies, we are witnessing a massive increase in the number of genomes in public repositories, with nearly a doubling per year in the Genomes OnLine Database (GOLD) [1, 2]. Reference genomes are important in both medical and environmental microbiology for capturing information on metabolic properties [3], phylogeny [4], evolution and diseases [5, 6], population genetics [7], functionality and biogeochemical cycles [8], interactions [9] and to establish links between genomes and functionality of cells in organisms [10]. In fact, obtaining good and relevant reference genomes is crucial for current advances in many, if not all, branches of biological research [11].

Prokaryotes dominate the biosphere in the context of abundance and diversity [12] and hold key roles in biogeochemical processes essential to all life [13]. However, only a small fraction of the bacterial diversity (<1%) can be isolated and cultivated in a standardized fashion [14]. Therefore, strategies for recovering genomes from samples without the need for cultivation have emerged as important complements to traditional microbiological techniques. In the single amplified genome (SAG) strategy, genomes of individual cells are sequenced. The first step comprises partitioning of the cells [15-17] using techniques such as fluorescent activated cell sorting (FACS) [18, 19] or microfluidics [20]. The next step involves cell lysis and whole genome amplification (WGA) for which three methods are most commonly used; PCR-based (e.g. degenerate oligonucleotide primed PCR (DOP-PCR)), isothermal (e.g. multiple displacement amplification (MDA)) or hybrid methods (e.g. multiple annealing and looping based amplification cycles (MALBAC)) [21] before applying shotgun sequencing and genome assembly [20, 22].

Genomes can also be recovered from metagenomes by assembling short shotgun reads into longer contigs which are then clustered into groups, or bins, of contigs derived from the same organism, through a process called binning. The resulting bins are quality filtered for contamination and completeness and the approved bins are sometimes referred to as metagenome assembled genomes (MAGs) [23]. This approach has been used for some time [24], but a fairly recent development is to perform the binning using a combination of sequence composition and differential abundance information [25-28]. Whereas it is possible to use as few as two samples for utilizing differential abundance information, the quality of the binning results can be greatly improved by increasing the number of samples [26, 27].

Although both the SAG and the MAG approach have proven powerful and contributed greatly to our understanding of the physiology and evolution of organisms [23, 29-32], a number of challenges are associated with each approach. SAG sequencing is demanding in terms of instrumentation and staff [33]. Starting with only one genome copy makes DNA amplification necessary but difficult, which often results in highly uneven coverage depth and some regions being completely missing from the sequencing output [21, 34]. Furthermore, cell dispersion, which might be necessary when cells are attached to particles or have formed biofilms, can be problematic and hinder genome recovery from some single-cells [35]. Obtaining a large number of high quality MAGs, on the other hand, requires extensive sequencing and ideally a large number of samples that to some degree share the same organisms in different abundances [27]. The quality of the MAGs is also highly dependent on the quality of the metagenome assembly; short contigs are not considered by most binning algorithms since their coverage and composition information contain too much noise [27, 36, 37]. Another limitation is the computational demands, which normally exceed those for SAG assembly [36]. Also, due to intraspecies genetic variation in the community, genomes recovered from metagenomic data often represent a population of closely related organisms (i.e. strains) rather than an individual organism [36].

Studies have successfully combined the SAG and MAG approaches to reach conclusions about organisms and ecosystems [38, 39]. The approaches have also been combined to methodologically improve either the quality of the single-cell assemblies [40] or the metagenome binning performance [41]. However, with the exception of a study that focused on a single phylum and that did not use abundance patterns over multiple samples for the MAG construction [38] the performance of the two approaches have to our knowledge not been thoroughly compared. The aim of this study was to do a comprehensive comparison between the SAG and MAG approaches for recovering prokaryotic genomes. We investigated SAGs and MAGs from bacterioplankton collected in the Baltic Sea Proper, where recent analyses have provided a detailed picture of the spatiotemporal distribution of microbial populations [23, 42-44] and metabolic processes [45]. Thus, this ecosystem is well suited for comparing different methodologies for investigating the genomic content and functional potential of dominant bacterial populations.

## Results

### Overview of SAGs and MAGs

In order to compare single-amplified genomes with metagenome-assembled genomes from the same environment, we generated SAGs from the Linnaeus Microbial Observatory (LMO), located 11 km off the coast of Sweden in the Baltic Sea, and compared them with MAGs generated earlier from the same station (Hugerth et al. 2015). We obtained 16 SAGs of a variety of taxa including *Bacteroidetes, Cyanobacteria, Alpha-* and *Gammaproteobacteria* (Table S1). These were compared to 83 MAGs from 30 phylogenetically distinct Baltic Sea clusters (BACLs) [23] (Fig. S1; Table S1). The SAGs ranged in size from 0.14 to 2.15 Mbp and MAGs from 0.59 to 2.98 Mbp (Table S1). The number of contigs in SAGs ranged from 80 to 712 with a maximum length of 107,141 bp, while the number of contigs in MAGs ranged from 60 to 951 with the longest being 181,472 bp (Table S1).

Using Mash [46] to cluster the 99 genomes from both approaches, seven of the 16 SAGs were placed together with 24 of the MAGs into six clusters (i.e. each of these SAGs matching 1-14 MAGs and each of these MAGs matching 1-2 SAGs; Table 1 and Fig. S1). This was in agreement with the clustering of MAGs in the analysis of Hugerth et al. [23]. These clusters belonged to a diverse set of bacterial taxa, representing the SAR86 and SAR92 clades *(Gammaproteobacteria), Flavobacteriaceae* (2 taxa) and *Cryomorphaceae (Bacteroidetes)* and *Rhodobacteraceae (Alphaproteobacteria)* (Table 1).

**Table 1.**
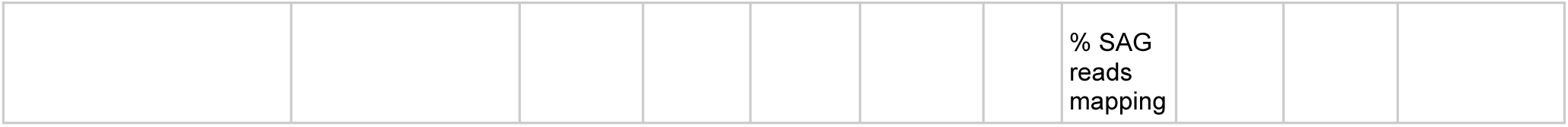
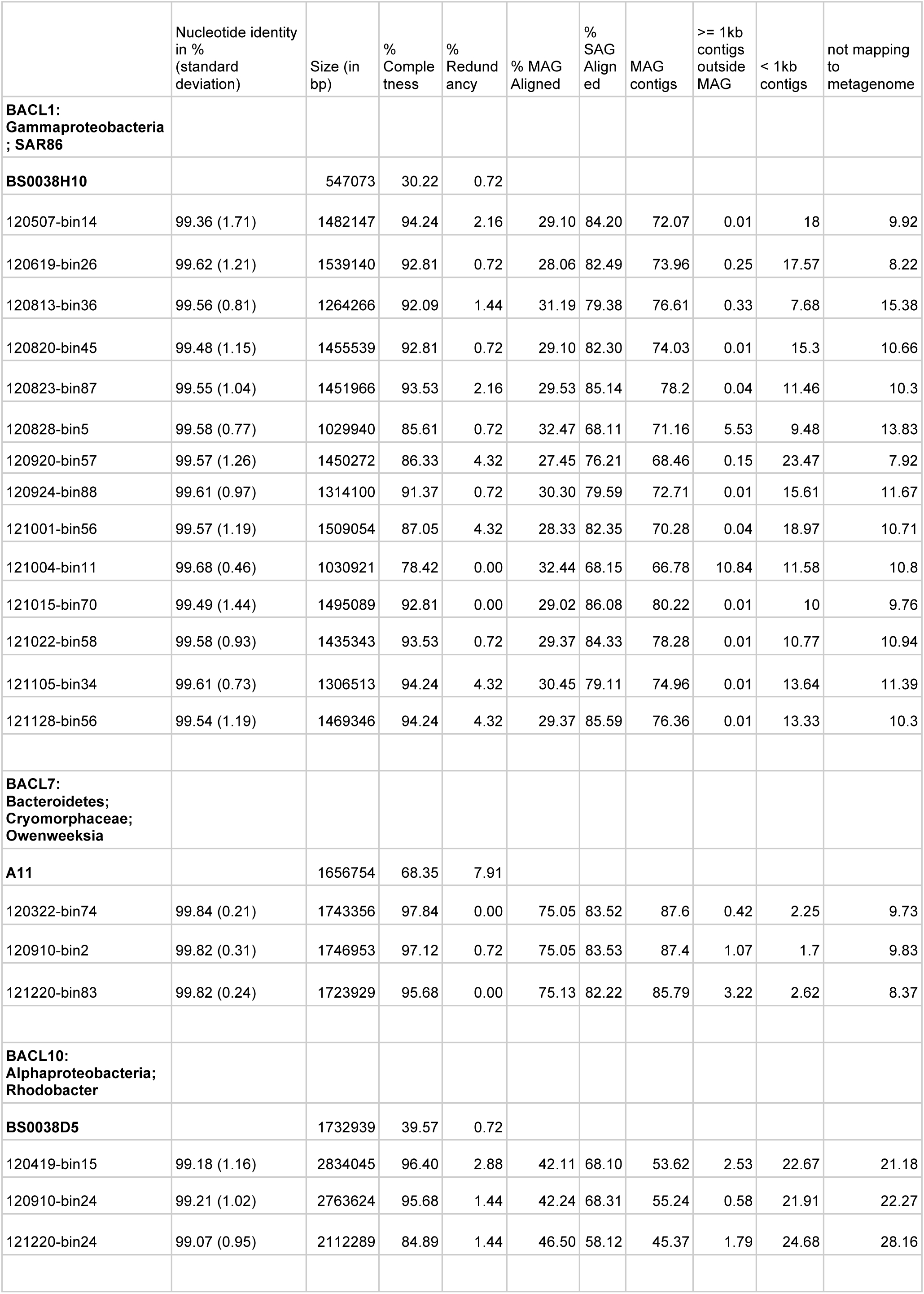
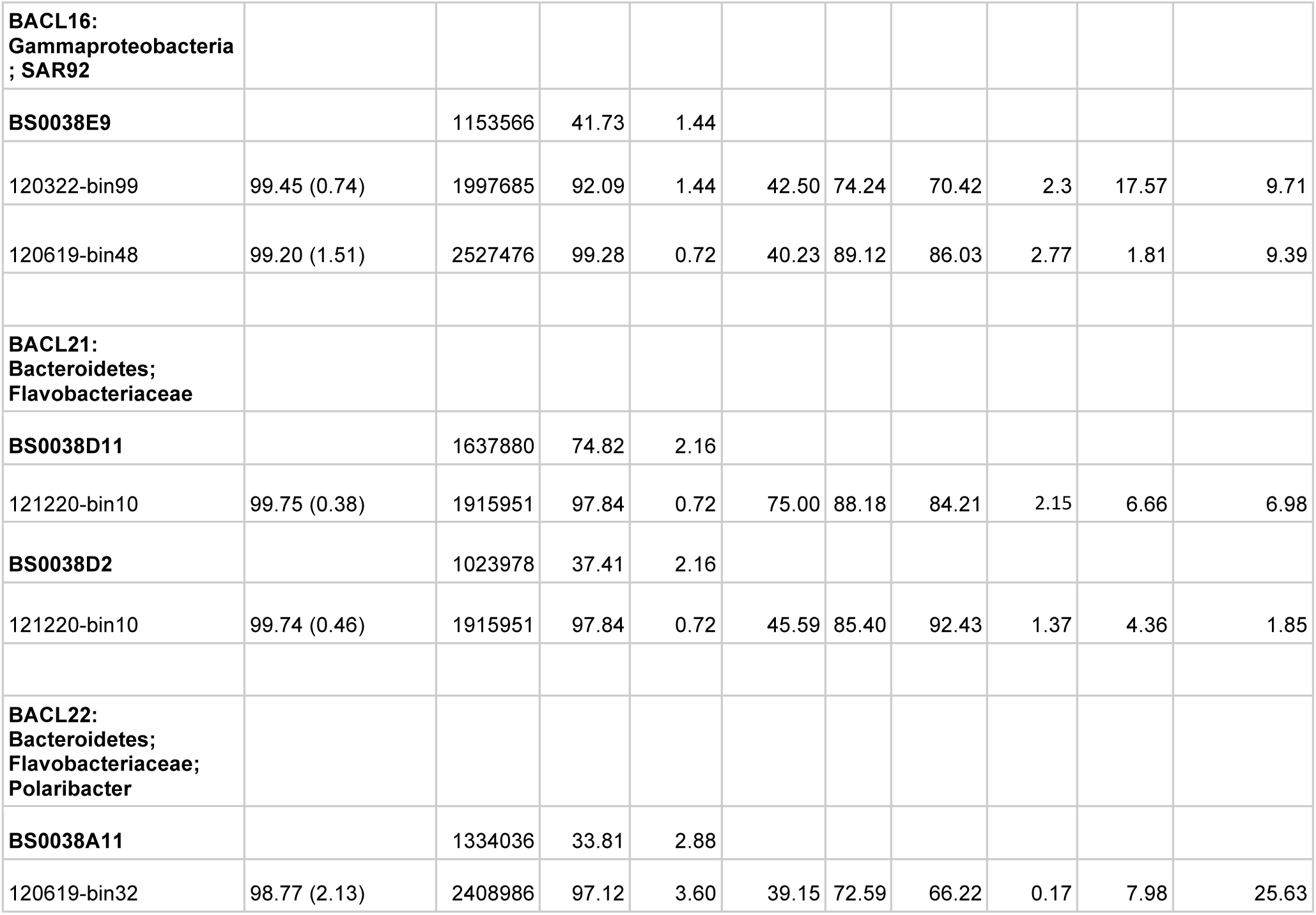
Overview of the matching SAGs and MAGs sorted by Baltic Sea cluster (BACL) number.

The seasonal dynamics of the clusters at the LMO station were determined in the original MAG study by metagenome samples covering a single year (2012) [23]. By comparing the 16S rRNA gene sequences from the genome clusters to 16S rRNA gene data from an amplicon-based high temporal-resolution study from the same station from the previous year (2011) [44], we observed five matches with a sequence identity of 100%. In these cases, the seasonal dynamics of the genome clusters and OTUs was similar between the years, with representatives abundant in spring and late autumn (2012), (BACL21, *Flavobacteriaceae*, OTU:000004 and BACL7, *Owenweeksia*, OTU:000021), spring and early summer (BACL16, SAR92 clade, OTU:000043), spring, summer and autumn (BACL10, *Rhodobacteraceae*, OTU:000011) and all year around (BACL1, SAR86 clade, OTU:000013) [23, 44] (Fig. S2). The contigs representing the genomes of BACL22 lacked the 16S rRNA gene sequence and were not included in the seasonality analysis.

### Alignment and Gene content

To verify the clustering and to achieve more detailed statistics, each SAG-MAG pair was aligned using MUMmer (Table 1). Across the genome regions showing homology between SAGs and MAGs, the within-cluster nucleotide identity was >98.7%. A larger fraction of the SAGs’ bases (average 78.9%) aligned compared to the MAGs’ (average 40.5%), in agreement with the SAGs being consistently smaller than the corresponding MAGs; 0.5-1.7 Mbp and 1.0-2.8 Mbp, respectively (Table S1) [23].

To further compare the SAGs and MAGs, the Anvi’o pangenomic workflow [47] was run on each cluster (Fig. 1, Table S2). This analysis showed that the completeness of the SAG genomes (average 46.6%) was lower than for MAG genomes (average 92.6%) (Table 1), as estimated by Anvi’o (by presence of 139 bacterial Single Copy Genes [SCGs]). Redundancy in gene content (measured as SCGs present more than once) showed no systematic difference between SAGs and MAGs (Table S2); it was highest in SAG A11 and in four MAGs of BACL1 (with 7.9% and 4.3% respectively).

**Figure 1.**
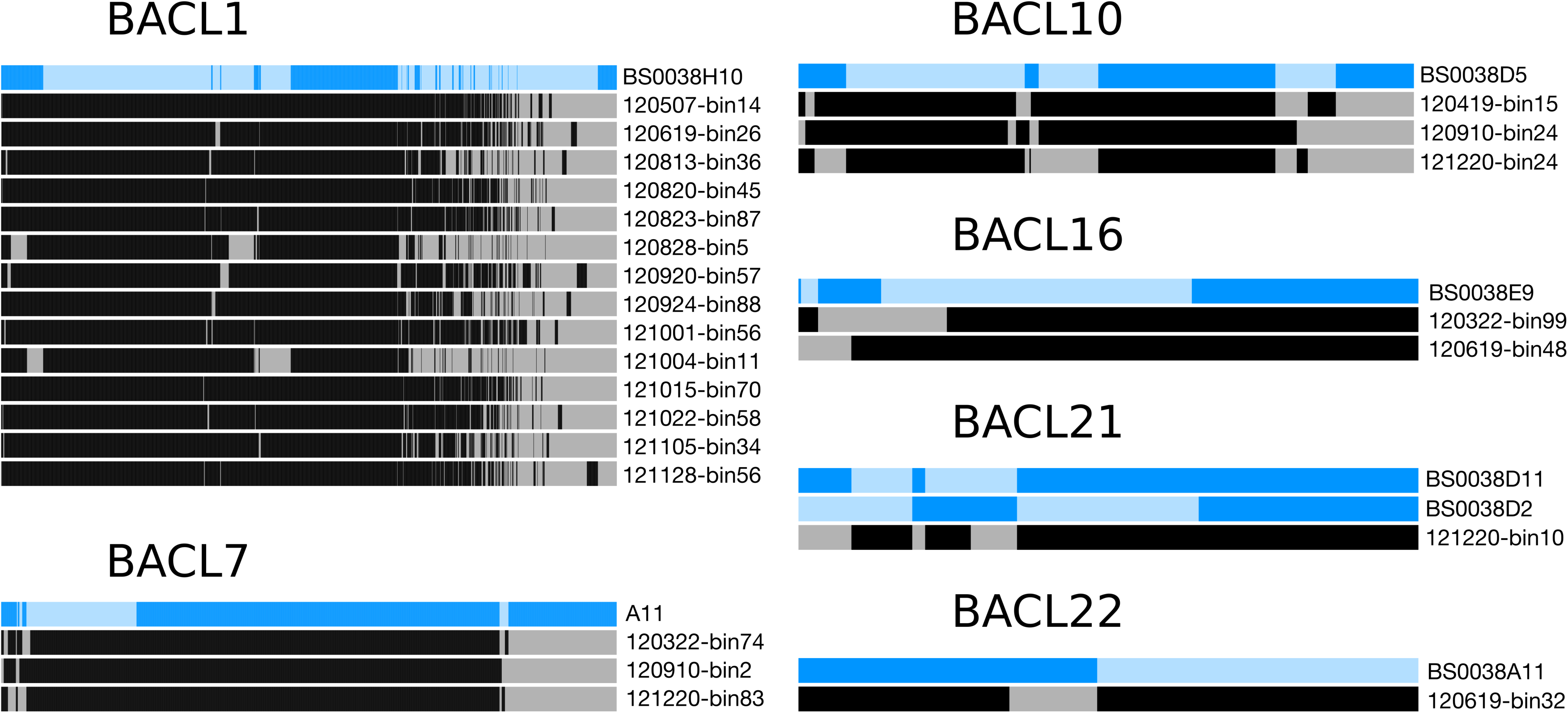
Gene homologs presence per genome cluster. Presence of gene homologs for each genome cluster by graphs produced by Anvi’o. Each horizontal bar represents one genome, where blue bars are single-amplified genomes and black and grey bars are metagenome-assembled genomes. Each vertical bar correspond to one gene homolog where a dark vertical bar indicate presence of the gene homologs and a lighter vertical bar indicate absence. The gene homologs are aligned between genomes within each genome cluster. The numbers assigned to the genome clusters corresponds to the original MAG BACLs used in [23].

There was a substantial range in gene content overlaps in different clusters (Fig. 1). For example, most MAGs in BACL1 contained a large set of genes (∽35% of genomes) missing in the corresponding SAG (BS0038H10), whereas the SAG in this cluster contained few genes (∽5% of genomes) not present in the MAGs. In contrast, in BACL7, similar portions of the genes (∽20% of genomes) were unique to the SAG or the MAGs. The case of BACL21 is particularly interesting since it contained two SAGs (the only cluster with more than one SAG) that differed substantially in size (1.0 Mb and 1.6Mb; Table 1). The two SAGs together covered nearly the entire gene content of the corresponding MAG (Fig 1).

### Analysis of functional gene data

Despite the differences in genome sizes, the distribution of functional genes as defined by Clusters of Orthologous Groups (COGs) of genes was consistent within SAG and MAG clusters. Hierarchical clustering based on counts of individual COGs reconstructed the genome clusters (Fig 2a), and broad functional COG categories were consistently distributed within clusters (Fig 2b). The distribution rather appeared to differ taxonomically. For instance the COG category “Amino acid metabolism and transport” was more abundant in the cluster BACL10 *(Rhodobacter)* compared to in other clusters. The *Flavobacteria* (BACL7, 21 and 22) showed elevated proportions of the functions “Cell wall/membrane/envelope-biogenesis” and “Translation”. “Lipid metabolism” was more frequent in the clusters of *SAR86* and *SAR92* clade (BACL1 and 16) compared to other clusters (Fig. 2b).

**Fig. 2.**
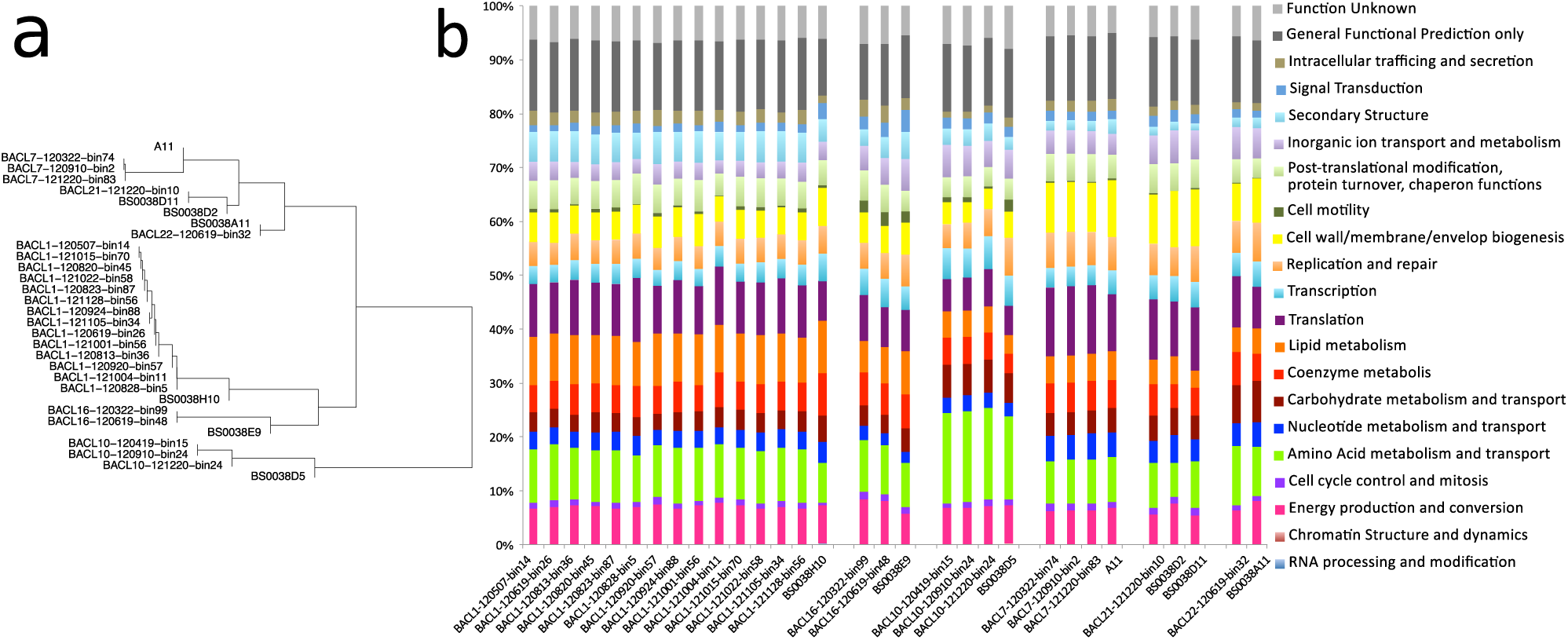
Distribution of functional categories per genome. **legend:** a) Hierarchical clustering of genomes based on their distributions of individual Clusters of Orthologous Groups (COGs) counts. b) Distribution of the COGs categories in the different genome cluster for MAGs and SAGs. Genomes, ordered according to genome clusters are displayed on the x-axis and on the y-axis is the percentage of COGs category in the cluster. The color represents the different broad categories used in the COG.

### Quantification of metagenome binning and assembly errors

Since the SAGs contained genome regions not present in the MAGs (on average 78.9% of SAG genomes aligned with the corresponding MAG genomes), we investigated potential reasons for these regions to be missing in the MAGs. Accordingly we determined the distribution of SAG sequencing reads mapping to different categories of metagenome contigs. This quantification showed that on average 73.9% of the SAG reads mapped to the contigs in their corresponding MAG (Fig 3a). Other metagenome contigs included in the binning, but that had hence ended up in other bins, recruited far fewer reads (average 1.42%)(Fig 3b). The length of DNA covered by SAG reads in these contigs divided by the length of DNA covered by SAG reads in the contigs that were subject to binning was on average 3.6% (Table S3). This estimate is valuable since it serves as an estimate of the false negative rate of the binning procedure. The remaining SAG reads were either mapping to small contigs (<1000 bp) not included in the binning because they were too short (<1000 bp) (average 12.44% of reads) or not mapping to metagenome contigs at all (12.20% of reads) (Fig 3c,d), and were hence rather reflecting insufficient metagenomic assembly.

**Fig. 3.**
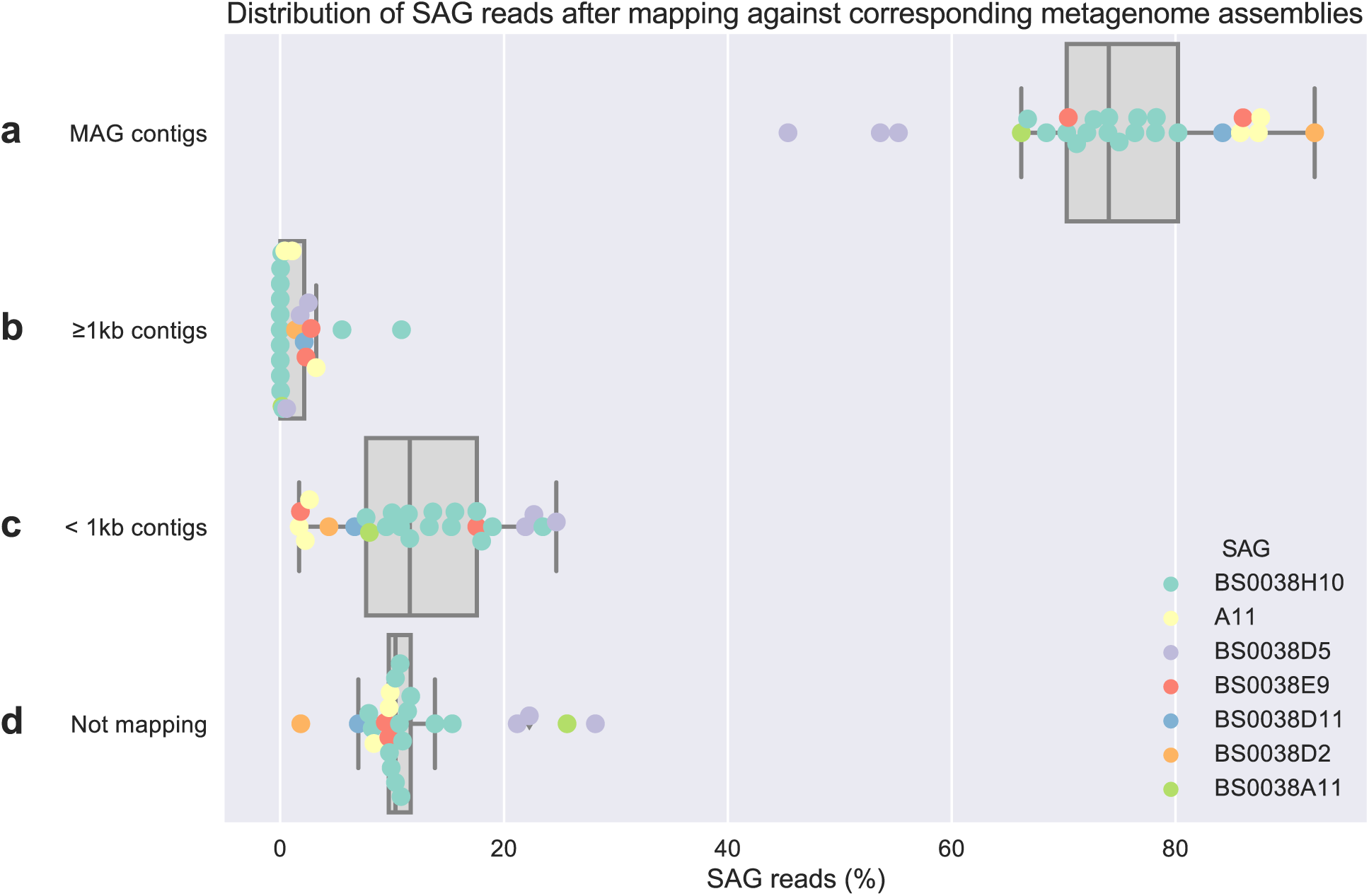
Distribution of SAG reads mapped against metagenome assemblies. **legend:** Boxplot of the distribution of SAG reads mapped against the corresponding metagenome assemblies where each individual data point is jittered on top of each box. All reads for each SAG was mapped against the assembly associated with each matching MAG and thus positioned in exactly one out of these four categories. Only contigs longer than one kilobase was included in the binning, which is the reason to use it as a divider here.

From visual inspection of genome alignments we discovered some MAG contigs aligning to multiple SAG contigs. This was caused by duplicated sequences within the SAGs, where the highest amount was found within the A11 SAG assembly. However this issue was resolved when using the most recent version of the assembly software (Spades version 3.10.1 instead of version 3.5), tested on the A11 SAG (data not shown).

## Discussion

In this study we compared the genome output from two state-of-the-art approaches for obtaining prokaryotic genomes representing abundant populations in the natural environment without cultivation. From a collection of SAGs and MAGs, we found an overlap in six clusters, representing a broad taxonomic range including *Gammaproteobacteria, Bacteroidetes* and *Alphaproteobacteria* that were nearly identical between the two groups, verifying previous results with high average nucleotide identity between SAGs and MAGs [38]. Due to seasonal recurrence of populations in the studied waters [23, 44], very high identity could be achieved despite samples used for SAG sequencing and MAG construction were collected one year apart. From the relative abundance of matching data on specific bacterial populations (OTUs), we conclude that both approaches provide genomic information on abundant taxa in the natural environment.

There are, however, differences between the two methods. When conducting sequencing of single-amplified genomes, one of the benefits is that the cells can be screened and the researcher can select particular cells to sequence, perhaps targeting a specific taxon or function. Furthermore, if one has only very few samples, producing SAGs may be preferable since the efficiency of the MAG approach improves with the number of samples [27]. Similarly, the MAG approach may have difficulties binning closely related strains that display similar abundance patterns, or alternatively closely related strains that display a wide variation in genetic content, since core-and strain-specific parts of the genomes may obtain different abundance patterns. SAGs also supply superior information on which nucleotide variants that co-occur within a genome (haplotypes), whereas for metagenomics, this information is limited to the read length, although computational approaches for haplotype reconstruction are emerging [48]. Nevertheless, metagenome-assembled genomes do recover a higher percentage of the genome compared to SAGs. Also, since reads from many individuals of each population are being sampled, population genomic analysis can be performed using the metagenome data [49-51], and additional information about the whole microbial community is obtained from the metagenome dataset, which is achieved with a more standard set of equipment compared to that needed for single-cell sequencing. Multiple samples are often beneficial for ecological investigations, making such projects suitable for MAG construction. Nevertheless, the fact that the genomes matched abundant OTUs with representatives from different taxonomic groups shows that both the SAG and the MAG approaches have a broad generality when applied to environmental samples.

### Size of SAGs compared to MAGs

The SAGs in this study were consistently smaller than the corresponding MAGs. This could be caused by either incomplete SAG assemblies or by metagenome contigs erroneously placed in MAGs by the binning algorithm. Looking closer at the case where two SAGs aligned to the same single MAG (i.e. BACL21), there was evidence that the smaller of the two SAGs (BS0038D2) was incomplete, i.e. it lacked a large fraction of genes that were shared by the second SAG and the MAG (Fig. 1). Our results therefore support the first explanation, which has been previously observed [52, 53]. Combining the sections included in the SAGs would also cover a higher proportion of the MAG than any of the two SAGs did individually (Fig. 1). Furthermore, MAGs showed a low level of redundancy, which would likely have been higher if MAGs contained a high degree of erroneously binned contigs. Finally, SAGs are also less complete than MAGs as estimated by SCGs.

The cause for incomplete SAGs could be either uneven or incomplete amplification of parts of the single genome copy [21]. The average sequencing depth was, however, one order of magnitude higher for the SAGs than the MAGs (Table S1) and in most cases the sequencing reads used were longer, which should be beneficial for assembly quality. We therefore conclude that the major causes for small SAGs are problems during the whole-genome amplification. These are well-known issues of the SAG approach, and attempts to improve this method are ongoing [54, 55] - for example, multiple SAGs from the same population can be sequenced for better coverage [34]. Even though the SAGs were smaller than MAGs, the analysis of COG categories within each matching SAG and MAG demonstrated that the two approaches capture the broad functional categories in a similar manner (Fig. 2).

### Metagenome assembly is the principal source for MAG incompleteness

Since the redundancy was low in all SAGs, it is a reasonable assumption that there was no contamination during the laboratory work and that all sequencing reads originated from the intended organism. With the caveats that the whole genome amplification of single cells generates uneven depth of coverage for different parts of the genome [21], and potential sequence variation between strains of the same population, this allowed us to investigate how well the MAGs accounted for the whole genome of that organism within the metagenome (Fig. 3). SAG reads mapping to contigs included in the corresponding MAG accounted for the largest fraction for all pairs of MAGs and SAGs, confirming the completeness of the MAGs (Fig 3a). In contrast, reads mapping to contigs which were longer than 1 kilobase but not included in the corresponding MAG (Fig 3b), likely indicated wrongly binned contigs or possibly indicated sequence variation between strains of the same population. In the MAG assembly, only contigs longer than 1 kilobase were used as input to the binning, because short contigs are difficult to cluster correctly [27]. A high percentage of reads in this category would indicate substantial false negative binning errors - this was not the case (Fig. 3b). Instead, the false negative rate of the binning was low - on average only 3.6% measured as number of genomic bases.

This can be compared against the two categories which both quantify incompleteness of the MAG caused by non-optimal metagenomic assembly. Either the SAG reads mapped to parts of the metagenome assembly with contigs that were not successfully assembled past the length cutoff of 1 kilobase (Fig. 3c), or the genomic region to which the SAG read corresponded did not assemble well enough for the read to map (Fig. 3d). Since these categories had substantially higher proportions of reads, our comparison indicated that the metagenome assembly process had a larger effect on the level of incompleteness of the MAGs than the binning process itself. Improvements of metagenome assembly strategies have recently been made [56, 57], possibly reducing the influence of this issue. Furthermore, the metagenome assembly probably resulted in short contigs due to either too low coverage or high intraspecific diversity within the sample. These results indicate that it is rewarding to invest in high coverage and to carefully optimize metagenome assembly if one wants to recover as complete MAGs as possible.

### No significant core genome enrichment in MAGs

A potential problem with binning using coverage variations over multiple samples is that strain-specific genes can have different abundance profiles than the core genome if multiple strains of the same species are present in the samples [27]. Therefore strain-specific and core genes are at risk of being placed into different bins and the use of single copy core genes as an estimate of completeness would result in an overly optimistic measure for the core genome bin. If any of the MAGs would be artificially core-genome-enriched in the binning procedure, we would expect a large fraction of the SAG reads, in particular those corresponding to the non-core genome, to map to the long contigs that were not in the MAGs. This was however not the case, as only a very small fraction was detected (Fig 3b). These findings indicate that core genome enrichment in the construction of MAGs is likely a smaller problem than the metagenome assembly.

## Conclusions and summary

Individual MAGs in this study were found to be larger and more complete than corresponding SAGs, although there is reason to believe that analysis of multiple SAGs from the same group of organisms could result in equal or higher completeness if jointly assembled. The false negative rate in the binning process was generally low and, instead, the metagenome assembly was found to be the largest hurdle to recover high quality MAGs. Single-cell-technology offers the possibility of genome recovery from a single sample whereas the reconstruction of MAGs often requires multiple samples. This on the other hand, provides ecological information based on the MAGs abundance variations across samples. The strong agreement between the SAG and MAG methodologies emphasizes that both are accurate and that the choice of approach should depend on the research questions and on available resources.

## Material and Methods

### Generation of MAGs

The MAGs used in the current study were obtained as previously described in Hugerth et al. [23]. Briefly, bacterial community DNA for MAG construction was obtained from surface water (2 m) collected in the Baltic Sea on 37 time points between March and December 2012 at the Linnaeus Microbial Observatory (LMO) located ∽11 km offshore Kârehamn, Sweden (56°55’.51.24 “ N 17°3’38.52 “ E). Library preparation of the bacterial community DNA were performed with the Rubicon ThruPlex kit (Rubicon Genomics, Ann Arbor, Michigan, USA) according to the instructions of the manufacturer and finished libraries were sequenced on a HiSeq 2500 (Illumina Inc., San Diego, CA, USA) with paired-end reads of 2 x 100 bp at SciLifeLab/NGI (Solna, Sweden). On average 31.9 million paired-end reads per sample were generated.

Quality controlled reads were assembled separately for each sample using a combination of Ray 2.1 (Ray Meta) [58] and 454 Life Science’;s software Newbler (v.29; Roche, Basel, Switzerland). Bowtie2 [59] was used to map all quality controlled reads for each sample against the contigs. Contigs from each sample were then binned using CONCOCT [27], an algorithm that clusters contigs into genomes across multiple samples, dependent on sample coverage and sequence composition using Gaussian mixture models. Bins were evaluated with a set of 36 single copy genes presented in [27] and approved if they contained at least 30 unique SCGs with a maximum of 2 in more than a single copy. Bins meeting these criteria were considered MAGs. Two MAGs from different samples could correspond to the same organism, and therefore, the 83 MAGs were clustered using MUMmer [60] into 30 Baltic Sea clusters (BACL). Functional analysis of each BACL was made with the PROKKA pipeline (v.1.7) [61] and extended with annotation for COG categories [62]. Taxonomic assignment for each MAG was firstly done with Phylosift [63] and then complemented with complete or partial 16S rRNA genes identified in the MAGs with webMGA [64].

### SAG sampling and Single cell sorting

Samples for SAGs from the Baltic Sea were collected 13th of May 2013 at the Linnaeus Microbial Observatory and cryopreserved in 1xTE, 5% glycerol (final concentration) before arriving to the Microbial Single Cell Genomics facility, SciLifeLab, Uppsala University. Prior to sorting, the cryopreserved samples were thawed, diluted, before being stained with 1x (final concentration) SYBR Green I (Life Technologies, CA USA) for approximately 30 minutes. The sorting was performed with a MoFlo Astrios EQ (Beckman Coulter, USA) cell sorter using a 488 nm laser for excitation, 70 μm nozzle, sheath pressure of 60 psi and 1.3 % sterile filtered NaCl as sheath fluid. Individual cells were deposited into 96-well plates (Bio-Rad, CA USA) containing 1 μl of 1x TE using a CyClone™ robotic arm and the most stringent single cell sort settings (single mode, 0.5 drop envelope). The sorter was triggered on forward scatter at a threshold of 0.08% and sort regions were set on SYBR Green I fluorescence detected at 513 nm using a 40 nm bandpass filter.

### Whole genome amplification using MDA with phi29

Deposited cells were lysed and neutralized followed by whole genome amplification using Phi29 and MDA as described by [18]. In short, the cells were incubated in an alkaline solution at RT for 5 minutes. Lysis reactions were neutralized by adding 1 μL neutralization buffer (Qiagen, Germany). MDA was performed using the RepliPHI™ Phi29 Reagent set (0.1 μg/μl, RH04210, Epicenter, WI USA) at 30°C for 16 h in 15 μ! reaction volumes with a final concentration of 1x reaction buffer, 0.4 mM dNTPs, 10 μΜ DTT, 5% DMSO, 50 μΜ hexamers with 3’- phosphorothioate modifications (IDT Integrated DNA Technologies, Iowa USA), 40 U Phi 29 enzyme; 0.5 μM SYTO13^®^ (Life Technologies, CA USA) and water. All reagents except SYTO13 were UV decontaminated at 2x 0.5 Joules in a Biolinker. The whole genome amplification was monitored in real time by detection of SYTO13 fluorescence every 15 minutes for 16 h using a Chromo4 real-time PCR instrument (Bio-Rad, CA USA). The single amplified genome DNA was stored at -20°C until further PCR screening, library preparation and Illumina sequencing.

### Screening of SAGs

Positive SAGs, defined by an early amplification curve well separated from negative controls as well as a positive PCR product targeting the 16S rRNA gene, were diluted 20-fold and screened using primer pair Bact_341 F: 5’- CCTACGGGNGGCWGCAG-3’and Bact_805 R: 5’- GACTACHVGGGTATCTAATCC-3’ [42]. The reactions were performed in 20 μl reaction volume with 2 U of Taq DNA Polymerase recombinant (Thermo Fisher Scientific, MA USA), 1x reaction buffer, 0.2 mM dNTPs, 2 mM MgCl2 and 0.25 μΜ of each primer. Following a 3 min denaturation at 95°C, targets were amplified for 35 cycles of 95°C for 30 s, 50°C for 30 s, 72°C for 60 s and a final 10 min extension at 72°C. PCR products were detected by an approximate 450 bp fragment on a 1.5 % agarose gel. The products were purified using the NucleoSpin Gel and PCR clean-up purification kit (Macherey-Nagel, Germany), quantified using the Quant-iT™ PicoGreen^®^ dsDNA assay kit (Invitrogen, MA USA) in a FLUOstar^®^ Omega microplate reader (BMG Labtech, Germany) and submitted for identification by Sanger sequencing at the Uppsala Genome Center.

### Illumina MiSeq sequencing

Altogether 15 SAGs were selected for genome sequencing. Twelve of these generated a 16S rRNA sequence identified by Sanger sequencing, and were selected to cover a broad range of phylogenetic groups. Three additional SAGs did not generate any 16S rRNA amplicons with the indicated primers, but were nevertheless selected to include also lineages not targeted by bacterial primers. The DNA content of the SAGs were quantified with the Quant-iT™ PicoGreen^®^ dsDNA assay kit and subsequently diluted to a concentration of 0.2 ng/μ! as recommended for the Nextera XT Library Preparation kit (Illumina, CA USA). Procedures were according to instructions from the manufacturer except that normalization was performed using the Kapa qPCR quantification method instead of bead normalization. In short, the Nextera XT uses an enzymatic step for fragmentation of DNA which enables small quantities of input DNA. The protocol involves a PCR amplification step where indexes and additional required nucleotide sequences are incorporated. After PCR cleanup, the library for each SAG was quantified and handed in for individual quality control at the SciLifeLab SNP&SEQ facility. The quality of the libraries was evaluated using the TapeStation from Agilent Technologies with the D1000 ScreenTape. The sequencing libraries were quantified by qPCR using the Library quantification kit for Illumina (KAPA Biosystems, MA USA) on a StepOnePlus instrument (Applied Biosystems, CA USA) and pooled in equal amounts prior to cluster generation and sequencing on a single MiSeq run with V3 chemistry and 2 x 300 bp mode.

One additional SAG (A11) from the same sample but from another sorted plate was purified using the NucleoSpin Tissue purification kit (Macherey-Nagel, Germany) and handed in directly to the SNPseq sequencing facility for preparation using the TruSeq Nano DNA library kit (Illumina, CA USA) and thereafter sequenced in another MiSeq V3 2 x 300 bp run.

### Data analysis of sequenced libraries

The global quality of raw and trimmed reads was checked using Fastqc 0.11 [65] and low quality data was removed together with adapters using Cutadapt 1.7 [66], requiring a minimal length of 75 nucleotides and using a quality of 30 as the threshold. The trimmed reads were assembled using the default values for Single Cell (-sc) with SPAdes 3.5 [67] and the parameter *carefull*, which, according to the documentation reduces the number of mismatches and short indels in contigs. The quality of each of the assemblies was assessed using the software QUAST 2.3 [68].

### Comparative genomics analyses

Mash version 1.0.1 [46] with 100,000 15-mers for each SAG and MAG was used to calculate pairwise distances between all genomes. Single linkage clustering was then performed using Scipy [69] and visualized using matplotlib [70] (Fig. S1). Clustering cutoff for each BACL was set at 0.1 (90% estimated similarity), and in each cluster containing a combination of MAGs and SAGs, they were pairwise aligned using the dnadiff tool from the Mummer suite version 3.23 [60]. Since Mash only gives an estimation of the nucleotide distance, we also subjected two additional clusters just over the 10% dissimilarity limit (BACL24 and BACL30) for alignment with MUMmer. Out of these, BACL30 resulted in the best alignment at 96.5% identity and alignment rate of the SAG at 53.7%. However, none of these two clusters were included in the comparison. The numbers assigned to the clusters corresponds to the original MAG BACLs used in [23].

Following the same procedure as [23], the SAGs were gene annotated using the PROKKA pipeline [61] and complemented with all significant (e-value < 0.00001) COG annotations using rpsblast from BLAST+ version 2.2.28+ [71]. The genomes were hierarchically clustered based on counts of COGs in the genomes using average-linkage clustering and pairwise genome distances calculated using poison dissimilarity [72] with the PoiClaClu package in *R* (www.r-project.org). Using the Anvi’o (Docker image with version 2.1.0) pangenomic workflow [47, 73] separately for each genome cluster, gene homologs were identified, visualized and estimates of completeness and redundancy were obtained. The summary statistics produced by Anvi’o are available in Table S2.

SAG reads corrected during the assembly process [67] that mapped to the SAG genome itself (minimum 99.55%) were mapped using Bowtie2 (version 2.2.6 with the --local argument) [59] against the assembled metagenome samples from which the MAGs were obtained. The resulting BAM-files were sorted using Samtools version 1.3 [74], duplicates were removed with Picard version 1.118 and the number of mapped reads per contig were counted (Fig. 3). Metagenomic contigs were divided into three groups: contigs included in the correct MAG, long (>=1 kb) contigs included in the binning but not belonging to the correct MAG and short (< 1 kb) contigs not included in the binning. Additionally there were those reads that did not map to the metagenome assembly at all. The counts were summarized and visualized using Pandas [75] and Seaborn [76].

Duplicated elements in the genomes were identified with BLASTN version 2.2.28+ [71] as alignments longer than 100bp between contigs longer than 1000bp and with 100% nucleotide identity. Reassembly of A11 was done using the corrected reads from existing assembly as input to Spades version 3.10.1 run in single cell mode.

### Seasonal occurrence of SAGs and MAGs

The majority of single-amplified genomes had 16S rRNA genes identified through Sanger sequencing as described previously. However, four SAGs (BS0038A02, BS0038A08, BS0038A11 and A11) lacked 16S rRNA gene sequence data and were therefore investigated with Barrnap (version *0.7)* [77]. In sample A11 the 16S rRNA gene was identified and taxonomically investigated using the SINA/SILVA database [78].

The three remaining samples BS0038A02, BS0038A08 and BS0038A11, were taxonomically annotated by keeping good quality contigs with a minimum length of 1kb and kmer coverage (provided by Spades) of at least 11.

Prodigal 2.6.1 [79] was then used to predict coding regions in the selected contigs and these contigs and predicted proteins were aligned against NCBI nucleotide and NCBI non-redundant database using BLAST (standalone BLAST + package version 2.2.30 [71]. 13 of the SAGs had a 16S rRNA gene which was individually blasted to 16S rRNA gene data from a field study at the LMO station [44] using online BLASTN [80]. The seasonal dynamics were then explored by comparing the matching SAG/MAG cluster from 2012 (i.e. BACLs from Hugerth et al 2015) to the corresponding OTU in 2011 (i.e. Lindh et al 2015).

## Additional Files

**Additional file 1: Fig. S1**. Hierarchical single-linkage clustering of SAGs and MAGs based on distances generated by MASH. Genome names starting with ‘BACL’ indicate MAGs and the number following indicates the Baltic Sea cluster. Leaves joined by nodes within a distance of 0.10 are grouped by colour of their leftmost branches.

**Additional file 2: Fig. S2.** Abundances over the year 2011 and 2012 for OTUs matching clusters of SAGs and MAGs. Redrawn from references Hugerth et al. and Lindh et al. [23, 44].

**Additional file 4: Table S1**. Assembly statistics and taxonomy for all MAGs and SAGs. For MAGs, coverage within the sample from where it was assembled.

**Additional file 5: Table S2.** Summary statistics as given by Anvi’o for all MAGs and SAGs found by both approaches.

**Additional file 6: Table S3.** Distribution of metagenome bases covered by SAG reads mapped against the corresponding metagenome assemblies. Estimate of false negative rate for the binning procedure.

## List of Abbreviations

**BACL:** Baltic Sea cluster

**COG:** Clusters of orthologous groups

**LMO:** Linnaeus Microbial Observatory

**MAG:** Metagenome-assembled genome

**Mbp:** Million base pairs

**OTU:** Operational taxonomic unit

**SAG:** Single-amplified genome

## Declarations

### Ethics approval and consent to participate

Not applicable

### Consent for publication

Not applicable

### Availability of data and materials

The single-amplified genome sequence dataset generated during the current study is available in the EMBL-EBI European Nucleotide Archive repository, under the primary accession PRJEB21451. The metagenomic reads dataset analysed in the current study are previously published [23] and are available on the sequence read archive under the accession SRP058493, https://www.ncbi.nlm.nih.gov/bioproiect/PRINA273799.

### Authors’ contributions

A.F.A. and J. P. conceived the study; J.A., C.M.G.K., A.F.A., and J. P. designed research; J.A., C.M.G.K., A.-M.D., C.B., F.H., M.V.L., L.W.H., T.J.G.E., S.B. performed research; J.A., C.M.G.K., A.F.A, and J.P. analyzed data; J.A., C.M.G.K., A.F.A., and J. P. wrote the paper.

## Acknowledgements

We thankfully acknowledge Anders Mânsson and Kristofer Bergström for their sampling at sea, and Sabina Arnautovic skilful support in the laboratory. We thank the National Genomics Infrastructure sequencing platforms at the Science for Life Laboratory at Uppsala University, a national infrastructure supported by the Swedish Research Council (VR-RFI) and the Knut and Alice Wallenberg Foundation. We’d also like to think the SciLifeLab Microbial Single Cell Genomics Facility at Uppsala University where the single cell genomics efforts were carried out. Computational resources, including support, were supplied to through UPPMAX: Uppsala Multidisciplinary Center for Advanced Computational Science, for which we are very grateful.

